# Off-the-shelf NIR-I fluorophores as ready-to-use NIR-II probes: screening and in vivo validation

**DOI:** 10.64898/2026.08.11.744199

**Authors:** Marie-Lynn Al-Hawat, Marc K. Saba-El-Leil, Simon Matoori

**Affiliations:** Faculté de Pharmacie, Université de Montréal, Montreal, QC H3T 1J4, Canada; Institute for Research in Immunology and Cancer (IRIC), Montréal, Quebec, Canada

**Keywords:** NIR-II fluorescence imaging, commercial fluorophores, cyanine dyes, DiR, liposomes, in vivo imaging

## Abstract

Fluorescence imaging in the second near-infrared window (NIR-II, 950-1700 nm) offers reduced scattering, lower autofluorescence, and deeper tissue penetration than NIR-I imaging, but its adoption is limited by the need for custom-synthesized fluorophores. Here, we identify commercially available dyes that exhibit usable NIR-II emission. Eleven visible, far-red, and NIR-I fluorophores were screened under twelve acquisition configurations combining 670, 760, and 808 nm excitation with band-pass (950 nm, 1400 nm) or long-pass (1000 nm, 1250 nm) emission filters. Output varied markedly with fluorophore identity and excitation/emission configuration. Among hydrophobic dyes, DiR exhibited strong emission across almost all excitation and emission filters. Among hydrophilic dyes, strong NIR-II fluorescence was observed for IRDye 680RD (excitation at 670 nm), sulfo-cyanine 7 (excitation at 670 nm and 760 nm), and indocyanine green (excitation at 808 nm). DiR showed a linear concentration-response under 760 nm excitation with BP1400 detection. Upon encapsulation in PEGylated liposomes, strong NIR-II fluorescence was retained. In an *in vivo* study in mice, NIR-II resolved vasculature that NIR-I could not consistently delineate, and enabled pharmacokinetic analysis. Both windows returned similar *ex vivo* organ distributions. NIR-II imaging is therefore accessible using commercial off-the-shelf fluorophores, provided the dye is matched to the intended excitation/emission configuration.

## INTRODUCTION

Fluorescence imaging in the second near-infrared window (NIR-II, 950-1700 nm) is emerging as a promising optical modality for biomedical research. Relative to visible and NIR-I (700-900 nm) imaging, photon scattering is markedly reduced, tissue autofluorescence is lower, and light penetrates deeper into tissue.^1–3^ The resulting increase in signal-to-background ratio enables sharper visualization of deep-seated vasculature and other structures at improved spatial resolution of probe biodistribution and longitudinal monitoring *in vivo*.^4,5^

Considerable synthetic effort has therefore been devoted to dedicated NIR-II fluorophores. Donor-acceptor-donor (D-A-D) small molecules, polymethine cyanines, aggregation-induced emission (AIE) luminogens, and semiconducting polymer nanoparticles have all been designed to obtain emission wavelengths beyond 1000 nm while improving molecular brightness, photostability, and *in vivo* imaging performance.^1,4,6,7^ These probes perform well, but most require multistep synthesis, extensive purification, or specialized expertise, which limits their accessibility and routine implementation outside dedicated chemistry laboratories.

Commercially available fluorophores, cyanine dyes in particular, constitute one of the largest families of probes used in biomedical research, owing to their low cost, availability, and well-characterized optical properties. Several isolated reports have shown that dyes originally developed for NIR-I imaging retain detectable emission in the NIR-II window, making them an attractive alternative to purpose-built NIR-II probes.^1,8,9^ These reports, however, have each examined single fluorophores rather than large libraries. We hypothesized that screening fluorophores from a library of commercial dyes would enable the identification of ready-to-use NIR-II fluorophores (Fig. 1). We therefore established a screening platform to assess a panel of commercially available fluorophores spanning visible, far-red, and NIR-I regions under twelve NIR-II acquisition configurations combining 670, 760, and 808 nm excitation with long-pass emission filters at 1000 nm and 1250 nm (LP1000, LP1250) and band-pass emission filters centered at 950 nm (BP950) and 1400 nm (BP1400). Due to its strong NIR-II fluorescence, DiR was selected for encapsulation into the membrane of PEGylated liposomes and evaluated by physicochemical characterization, *in vivo* imaging, and *ex vivo* biodistribution analysis.

**Figure 1.**
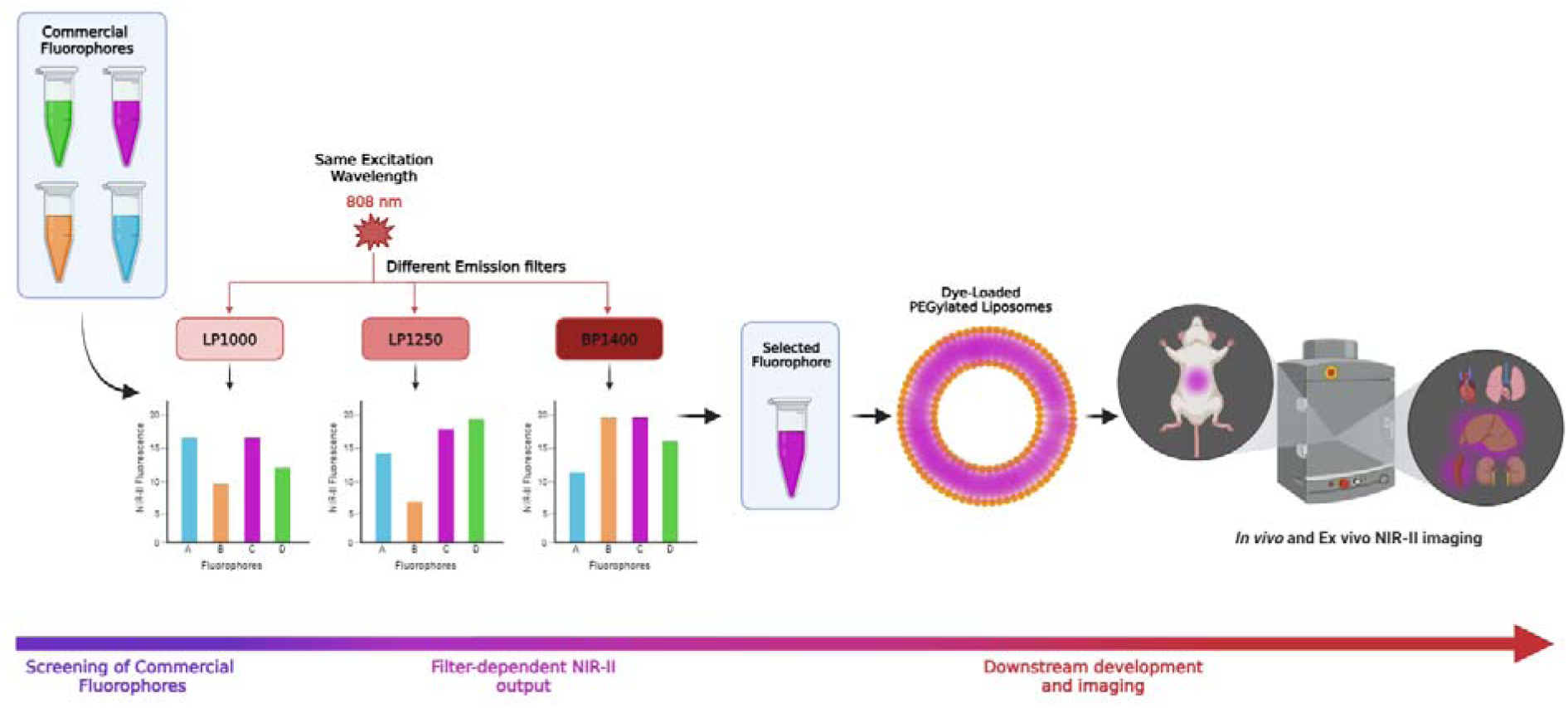
Concept and workflow for identifying off-the-shelf commercial fluorophores for NIR-II imaging. A panel of commercially available fluorophores spanning the visible, far-red, and NIR-I regions was excited at a single wavelength and imaged through a series of long-pass and band-pass emission filters (BP950, LP1000, LP1250, BP1400). This empirical approach is warranted because NIR-II emission relies on the tail of the emission band rather than its maximum. Therefore, manufacturer-reported spectra in the visible, far-red, or NIR-I range alone generally do not predict which dyes remain detectable at NIR-II wavelengths. A fluorophore retaining measurable signal under the most stringent filter was selected for downstream validation: incorporation into PEGylated liposomes, followed by *in vivo* and *ex vivo* NIR-II imaging in mice.

## MATERIALS AND METHODS

### Materials

The lipophilic fluorescent dyes DiA, DiO, DiI, DiD, DiR and the hydrophilic Alexa Fluor 647, indocyanine green (ICG), Sulfo-Cyanine7 carboxylic acid (S7) and Sulfo-Cyanine7.5 NHS Ester (S7.5) were bought from Lumiprobe (Hunt Valley, MD, USA). IRDye 680RD NHS Ester and IRDye 800CW were obtained from LI-COR Biosciences (Lincoln, USA). 1,2-dipalmitoyl-sn-glycero-3-phosphocholine (DPPC), cholesterol and 1,2-distearoyl-sn-glycero-3-phosphoethanolamine-N-[methoxy(polyethylene glycol)-2000] (DSPE-PEG(2000)) were obtained from Lipoid GmbH (Ludwigshafen, Germany). HEPES, sodium chloride, sucrose and ethanol were obtained from Sigma-Aldrich (St. Louis, MO, USA). PBS 10X (311-012-CL) was obtained from Wisent Inc. (Saint-Bruno, QC, Canada).

## Methods

### Screen of fluorophore library

A SynIRgy NIR-II imaging system (Photon etc., Montreal, QC, Canada) equipped with three excitation laser sources at 670, 760, and 808 nm, multiple bandpass and longpass filters on filter wheels, and an InGaAs camera, was used to evaluate the fluorescence properties of DiO, DiA, DiI, DiD, DiR, S7, IRDye 680RD, AF647, IRDye 800CW, ICG and S7.5.

All fluorophores were prepared at an identical concentration of 0.0005 mg/mL. DiA, DiO, DiI, DiD and DiR were dissolved in ethanol, whereas S7, IRDye 680RD, AF647, ICG and S7.5 were dissolved in ultrapure water. DiR in ethanol was also prepared at concentrations ranging from 0 to 500 ng/mL. All samples were imaged under identical acquisition conditions. Fluorescence images were acquired using different excitation wavelengths (670, 760 and 808 nm), emission filters: BP950 (900-1000 nm), LP1000 (1000-1650 nm), LP1250 (1260-1650 nm), and BP1400 (1375-1425 nm), exposure times and solvent conditions to optimize the imaging protocol. Fluorescence images were analyzed using Physpec v2 software (Photon etc.). Manual drawings of regions of interests (ROIs) were made, and mean fluorescence intensities were recorded. The acquisition parameters providing the highest signal-to-background ratio were selected for subsequent experiments.

### Preparation of fluorescent liposomes

Fluorescent liposomes were prepared using the thin-film hydration method.^10–12^ Briefly, DPPC, cholesterol and DSPE-PEG(2000) were dissolved in chloroform at a molar ratio of 45.4:53.7:0.9 mol% (DPPC:cholesterol:DSPE-PEG(2000)). DiR was added at a nominal dye-to-lipid ratio of approximately 1.5 mol%. The lipids and DiR were dissolved in chloroform, and the organic solvent was removed under a gentle nitrogen stream, followed by vacuum desiccation for at least 24 h to obtain a dry lipid film. The lipid film was hydrated with 20 mM HEPES (314 mosm/L) containing 10% (w/v) sucrose (pH 7.3) at 52 °C, and the resulting suspension was homogenized by alternating sonication and vortex mixing. Liposomes were subsequently diluted and extruded through polycarbonate membranes (100 nm, followed by 80 nm) at 55 °C using LIPEX extruder (Northern Lipids Inc., Vancouver, Canada) to obtain a homogeneous nanoparticle suspension with a total lipid concentration of approximately 19.3 mM. Liposome size, polydispersity index (PDI), and zeta potential were determined by dynamic light scattering using a Zetasizer (Malvern Panalytical, Malvern, UK).

DiR-loaded liposomes (initial nominal DiR concentration: 0.297 mg/mL) were serially diluted using sucrose-containing isotonic HEPES buffer (pH 7.3). Samples were imaged using the optimized acquisition parameters. And ROI analysis was performed to quantify fluorescence intensity, and with correlated DiR sensitivity and linearity of detection was evaluated.

### Animals

Male CD-1 (Crl:CD1(ICR); strain 022CD1; Charles River Laboratories) male mice were maintained under standard conditions at the Institute for Research in Immunology and Cancer. Mice were housed under specific pathogen-free conditions at a temperature of 23±1 °C, 55±5 % humidity, and a 12 h light/dark cycle in filter-topped isolator cages with access to food and water ad libitum. All animal procedures were performed in accordance with local animal welfare committee of the University of Montreal (Comité de Déontologie en Expérimentation Animale, CDEA; Protocol No. 23-018) in agreement with regulations of the Canadian Council on Animal Care (CCAC).

Animals were purchased at 5 weeks and were pair housed on arrival. Studies commenced after an acclimatization for a period of 7 days. Mice were single housed and randomized based on body weights the day before the study and had abdomens shaved using clipper and epilation cream. On the day of the pharmacokinetics study, body weights were recorded and used to calculate volume of administration.

### *In vivo* fluorescence imaging

Using the lateral tail vein, DiR-loaded liposomes were administered intravenously at a dose of 50 mg/kg per mouse in a volume of 10 mL/kg. At 1, 2, 4, 6 and 24 h post-injection, whole-body fluorescence imaging was performed using the optimized NIR-I (670 nm excitation, BP785 emission filter) and NIR-II (760 nm excitation, BP1400 emission filter) imaging parameters. Images were analyzed using PHySpec v2 software (Photon etc.) by measuring the mean fluorescence intensity within predefined ROIs encompassing the liver-associated region and the right femoral blood vessels.

### Plasma fluorescence analysis

Blood collection: Blood samples were collected (35 μL each) from individual mice at selected time points 1 h, 2 h, 4 h, 6 h, and 24 h following intravenous administration of DiR-loaded liposomes. Blood was collected from tail tip by gentle massage using a Minivette POCT-K3 EDTA blood collection capillary (Sarstedt, Nümbrecht, Germany). Fresh blood was transferred to a 0.5 mL Eppendorf tube and centrifuged at 9615 x g at 4°C for 10 min. Following centrifugation, 10 μL of plasma fraction was transferred to fresh tube and frozen on dry ice before being stored at -80°C. Plasma fluorescence was measured using a monochromator-based plate reader (Spark Multimode Microplate Reader, Tecan, Ma nnedorf, Switzerland) using a 745 nm excitation and a 800 nm emission wavelength. Mean fluorescence intensity was quantified and plotted as a function of time.

### *Ex vivo* biodistribution

Animals were euthanized and major organs such as the heart, lungs, liver, spleen and kidneys were collected 24 hours after administration. Organs were rinsed briefly in PBS 1x before imaging under NIR-I conditions using 670 nm excitation and a BP785 emission filter, and NIR-II conditions using 760 nm excitation and a BP1400 emission filter. Fluorescence intensity was quantified by ROI analysis for each organ and compared between NIR-I and NIR-II imaging conditions.

### Statistical analysis

Data are presented as mean ± standard deviation (SD). Statistical analyses were performed using GraphPad Prism version 11 (GraphPad Software, San Diego, CA, USA). For the fluorophore-screening experiments, each excitation–emission configuration was analyzed independently by one-way analysis of variance (ANOVA), followed by Tukey’s post-hoc test. Differences were considered statistically significant at p < 0.05. Significance levels are indicated as follows: *p < 0.05, **p < 0.01, ***p < 0.001, and ****p < 0.0001.

## RESULTS AND DISCUSSION

### *In vitro* screening of commercial fluorophores for NIR-II imaging

A panel of eleven commercial fluorophores spanning the visible, far-red, and NIR-I regions (DiO, DiA, DiI, DiD, AF647, IRDye 680RD, DiR, S7, IRDye 800CW, ICG, and S7.5) was screened at 0.5 µg/mL under twelve NIR-II acquisition configurations combining 670, 760, and 808 nm excitation with BP950, LP1000, LP1250, and BP1400 emission filters (Figure 2A-L). NIR-II output varied markedly across both fluorophores and configurations. DiR was the only fluorophore to remain significantly above its solvent control in all twelve configurations and gave very high signals across all excitation and emission wavelengths.

**Figure 2.**
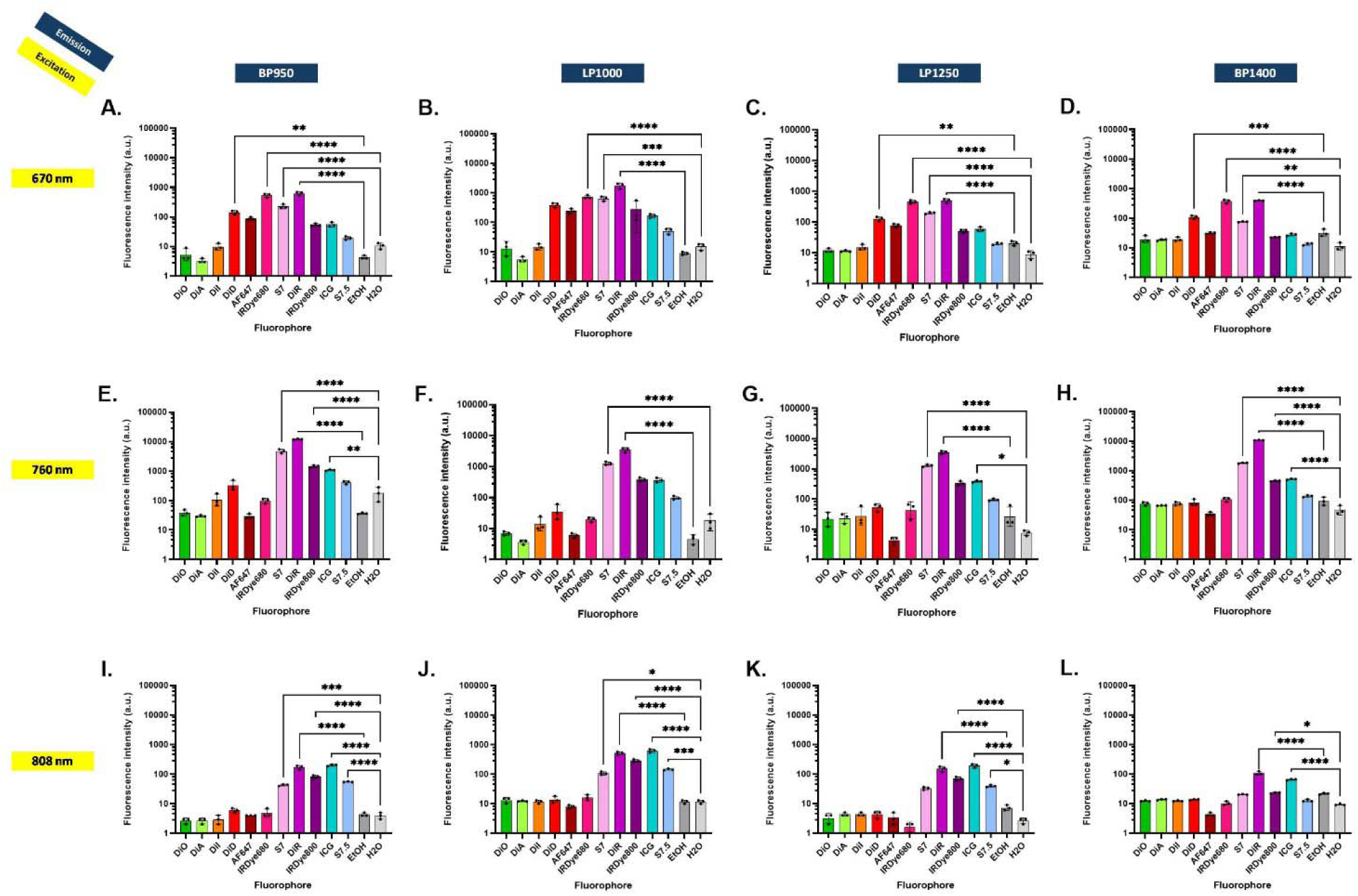
NIR-II fluorescence of commercially available fluorophores under different excitation–detection configurations. Fluorescence intensity of the screened fluorophores at a concentration of 0.5 µg/mL using 670 nm excitation with BP950 (A), LP1000 (B), LP1250 (C), and BP1400 (D) emission filters; 760 nm excitation with BP950 (E), LP1000 (F), LP1250 (G), and BP1400 (H) emission filters; and 808 nm excitation with BP950 (I), LP1000 (J), LP1250 (K), and BP1400 (L) emission filters. Mean fluorescence intensity was determined within manually defined regions of interest. Ethanol (EtOH) and ultrapure water (H O) were included as solvent controls. The y-axe is on a logarithmic scale. All results are mean ± SD (n = 3).

Under 670 nm excitation, IRDye 680RD and DiR produced significantly stronger fluorescence across all four emission windows than their respective negative controls (Figure 2A-D). Under 760 nm excitation, S7 and DiR both exhibited stronger fluorescence than their control across all four emission windows (Figure 2E-H). Under 808 nm excitation, ICG and DiR exhibited significantly higher signals than their control at all four emission wavelengths (Figure 2I-L). ICG has previously been identified as a NIR-II fluorophore through its long-wavelength emission tail, typically using 808 nm excitation and detection above 1000 nm^13^. The visible dyes DiO, DiA, and DiL as well as the far-red dye AF647 were indistinguishable from the controls throughout. IRDye 680RD fell off sharply under 760 and 808 nm excitation despite its strong performance at 670 nm. This finding demonstrates that selecting excitation wavelength close to the absorption maximum seems important to enable NIR-II emission. The NIR-II emission of IRDye 680RD also demonstrates that detectable NIR-II emission was not confined to the most red-shifted dyes. Because NIR-II detection relies on the tail of the emission band rather than its maximum^14,15^, signal depends on the extent of that tail within the detection window, on the extinction coefficient at the chosen excitation wavelength, and on the quantum yield in the relevant environment^1^. Solvent is a further contributor here, as the lipophilic dyes were dissolved in ethanol and the water-soluble dyes in aqueous buffer, conditions under which cyanine aggregation and self-quenching differ^13,16^. Of note is that dyes were compared at equal mass rather than equal molar concentration (SI Table S1).

To assess linearity and potential saturation, a calibration curve was generated from serial dilutions of free DiR under 760 nm excitation and BP1400 emission (Figure 3, SI Figure S2). Fluorescence intensity was linear with concentration across the range tested (R² = 0.9923). This linear response, combined with the high NIR-II signal and the established use as liposomal membrane dye of DiR^17^ supported its selection for incorporation into PEGylated liposomes.

**Figure 3.**
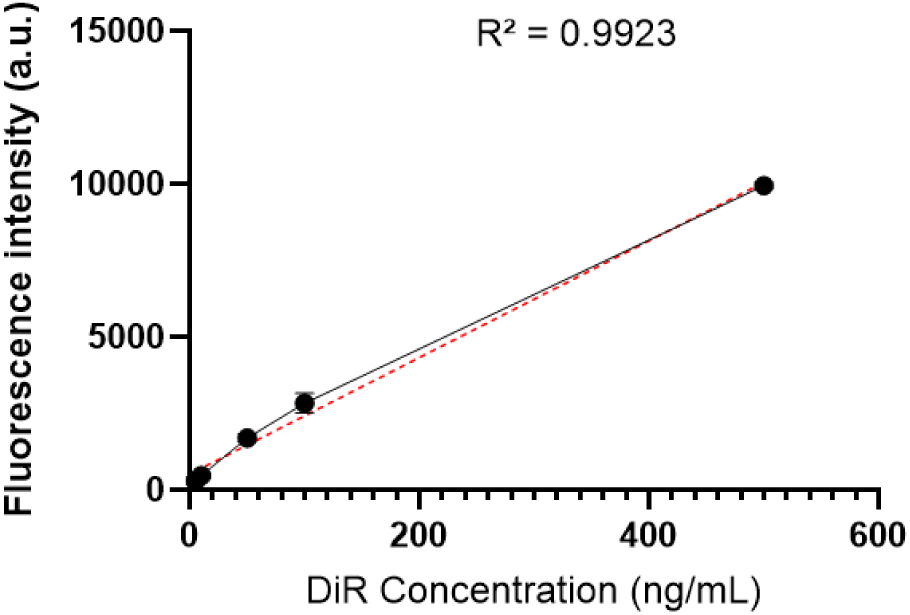
Concentration-dependent NIR-II fluorescence response of free DiR. Fluorescence intensity of free DiR solutions at concentrations ranging from 0 to 500 ng/mL using 760 nm excitation and a BP1400 emission filter. Mean fluorescence intensity was determined within manually defined regions of interest and plotted as a function of DiR concentration. The red dashed line represents the linear regression with a coefficient of determination of R² = 0.9923. All results are mean ± SD (n = 3).

### Characterization of DiR-loaded liposomes

DiR was encapsulated in the membrane of PEGylated liposomes. DiR-loaded liposomes were first characterized physicochemically (Table 2). They exhibited a mean hydrodynamic diameter of 92 nm with a polydispersity index of 0.100, indicating a narrowly distributed and homogeneous population within a range adapted for intravenous administration.^18,19^ The moderately negative zeta potential of -13.5 mV is consistent with similar PEGylated liposomal formulations.^19^

**Table 1.**
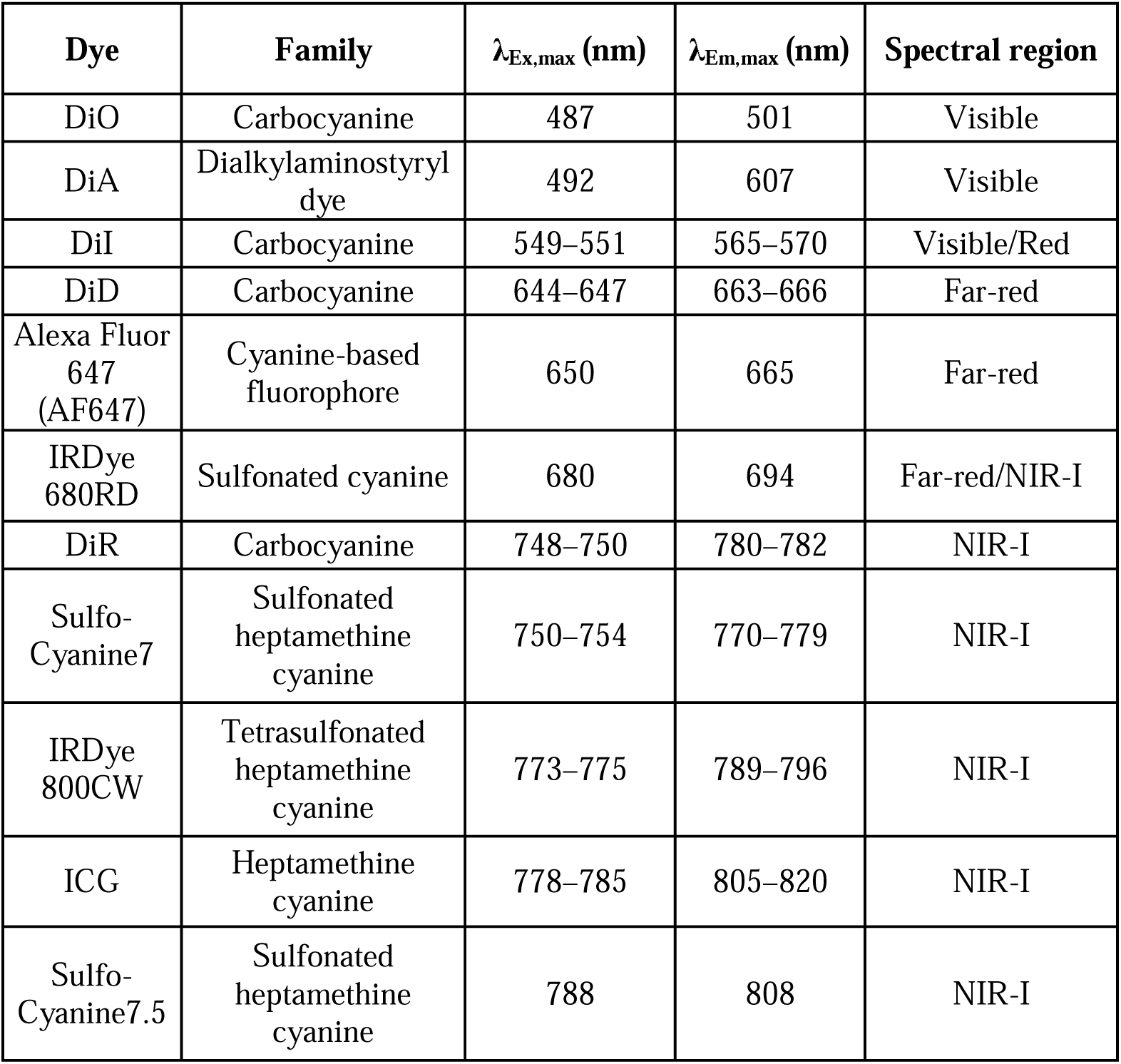
Spectral characteristics of the commercial fluorophores included in the screening panel. Excitation maxima (λ_Ex,max_), emission maxima (λ_Em,max_), fluorophore families, and corresponding spectral regions are indicated as reported by manufacturer.

| Dye | Family | $\lambda_{\text{Ex,max}}$ (nm) | $\lambda_{\text{Em,max}}$ (nm) | Spectral region |
| --- | --- | --- | --- | --- |
| DiO | Carbocyanine | 487 | 501 | Visible |
| DiA | Dialkylaminostyryl dye | 492 | 607 | Visible |
| DiI | Carbocyanine | 549–551 | 565–570 | Visible/Red |
| DiD | Carbocyanine | 644–647 | 663–666 | Far-red |
| Alexa Fluor 647 (AF647) | Cyanine-based fluorophore | 650 | 665 | Far-red |
| IRDye 680RD | Sulfonated cyanine | 680 | 694 | Far-red/NIR-I |
| DiR | Carbocyanine | 748–750 | 780–782 | NIR-I |
| Sulfo-Cyanine7 | Sulfonated heptamethine cyanine | 750–754 | 770–779 | NIR-I |
| IRDye 800CW | Tetrasulfonated heptamethine cyanine | 773–775 | 789–796 | NIR-I |
| ICG | Heptamethine cyanine | 778–785 | 805–820 | NIR-I |
| Sulfo-Cyanine7.5 | Sulfonated heptamethine cyanine | 788 | 808 | NIR-I |

**Table 2.** Physicochemical characteristics of DiR-loaded PEGylated liposomes. Z-average hydrodynamic diameter, peak diameter by intensity, polydispersity index (PDI), and zeta potential were determined by dynamic light scattering.

| Parameter | Value |
| --- | --- |
| Z-average diameter (nm) | 91.68 |
| Peak diameter by intensity (nm) | 102.4 |
| Polydispersity index (PDI) | 0.100 |
| Zeta potential (mV) | -13.53 |

To determine whether DiR retained its NIR-II properties after incorporation into the liposomal membrane, serial dilutions of the liposomal formulation were imaged under the same conditions used for the screen. Fluorescence decreased progressively with dilution and remained readily detectable across the range tested (Figure 4A). Over the full series, however, intensity deviated from linearity and approached a plateau at the highest concentrations (Figure 4B); regression was therefore restricted to the initial linear portion (Figure 4C). The plateau may reflect self-quenching of DiR concentrated within the lipid bilayer, detector saturation, or an inner-filter effect^20^. Quantitative NIR-II measurements of this formulation should be restricted to the experimentally established linear range where DiR-loaded liposomes can be reliably quantified under NIR-II imaging conditions.

**Figure 4.**
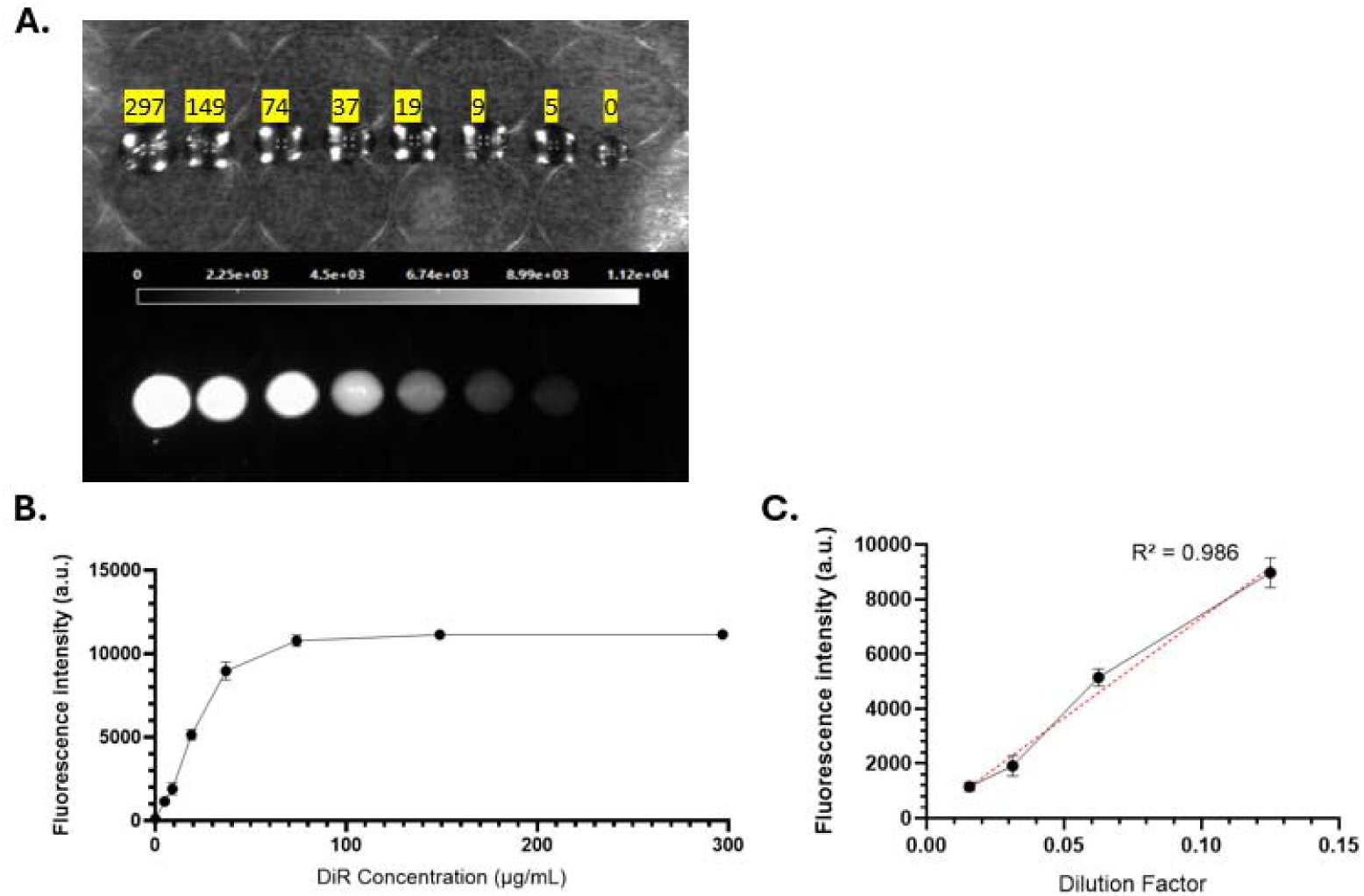
NIR-II fluorescence response and linear dynamic range of DiR-loaded liposomes. Representative broadband and NIR-II fluorescence images of DiR-loaded liposomes at nominal DiR concentrations of 0, 5, 9, 19, 37, 74, 149, and 297 µg/mL acquired using 760 nm excitation and a BP1400 emission filter (A). Mean fluorescence intensity plotted as a function of Di concentration over the complete dilution series (B). Linear regression of the initial linear portion of the dilution curve (C). The red dashed line represents the linear regression, with a coefficient of determination of R² = 0.986. All results are mean ± SD (n = 3).

### *In vivo* imaging of NIR-I and NIR-II fluorescence imaging of DiR-loaded liposomes

DiR-loaded liposomes were administered intravenously to mice and imaged for 24 h under matched NIR-I (670 nm excitation, BP785) and NIR-II (760 nm excitation, BP1400) conditions. Both modalities identified the liver as the principal site of accumulation over 24 h (Figure 5A), but NIR-II images showed reduced diffuse background and clearer delineation of major vessels, particularly during the first hours after injection (Figure 5B).

**Figure 5.**
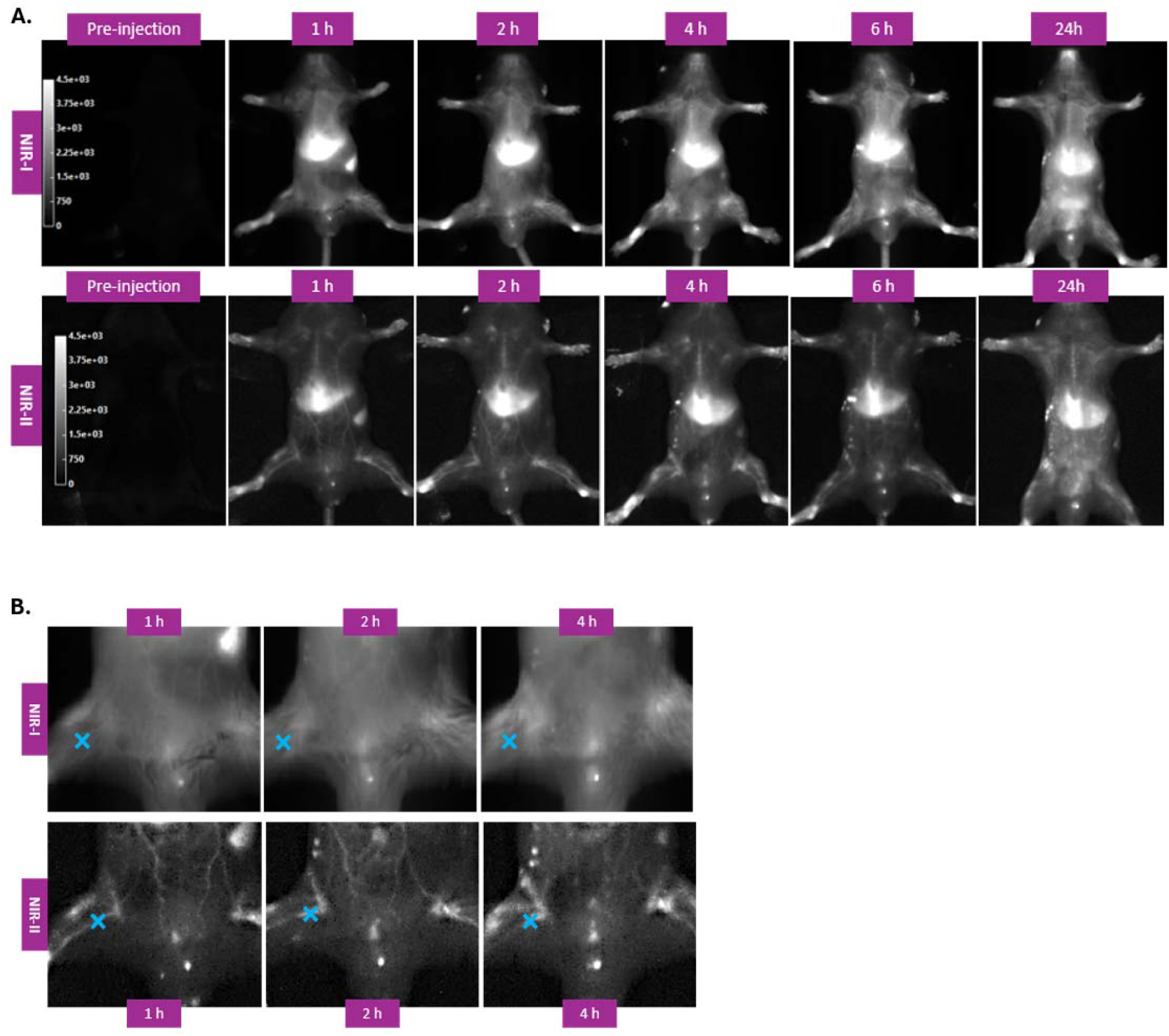
*In vivo* NIR-I and NIR-II fluorescence imaging following intravenous administration of DiR-loaded liposomes. Representative longitudinal whole-body fluorescence images acquired before injection and at 1, 2, 4, 6, and 24 h after intravenous administration of DiR-loaded liposomes (A). NIR-I images acquired using 670 nm excitation and a BP785 emission filter (upper row), and NIR-II images acquired using 760 nm excitation and a BP1400 emission filter (lower row). Images are displayed using identical intensity scales within each imaging modality. Magnified images of the abdominal region were acquired at 1, 2, and 4 h under NIR-I conditions (upper row) and NIR-II conditions (lower row) (B). Blue crosses indicate the right femoral blood vessel selected for quantitative fluorescence analysis. Representative images from one mouse are shown.

This difference was reflected in the quantitative time courses. Under NIR-II, liver-associated fluorescence rose sharply within the first hour and then plateaued and fluorescence in a major vessel peaked early and declined thereafter (Figure 6B). This pattern exhibits the typical biodistribution pattern of circulating liposomes.^21^ Under NIR-I, the hepatic accumulation were also observed, but the vascular ROI showed no distinct temporal trend (Figure 6A), as higher tissue autofluorescence reduced the signal-to-noise ratio. NIR-II thus yielded a more interpretable vascular pharmacokinetics profile despite lower absolute signal intensity, consistent with the reduced scattering and autofluorescence expected at longer wavelengths.^1,2^ Plasma fluorescence determined using a plate reader decreased steadily over the same period (Figure 6C), indicating progressive clearance of circulating liposomes. That the vascular time course of NIR-II imaging paralleled the decline in plasma fluorescence measured by a plate reader suggests that a vascular ROI could serve as a non-invasive surrogate for circulating liposome concentration, allowing blood-compartment kinetics to be followed at high temporal resolution without the need for invasive repeated blood sampling. Rapid hepatic uptake is consistent with the established pharmacokinetics of PEGylated liposomes, which are cleared from circulation primarily by hepatic Kupffer cells and splenic macrophages.^18,19,22^

**Figure 6.**
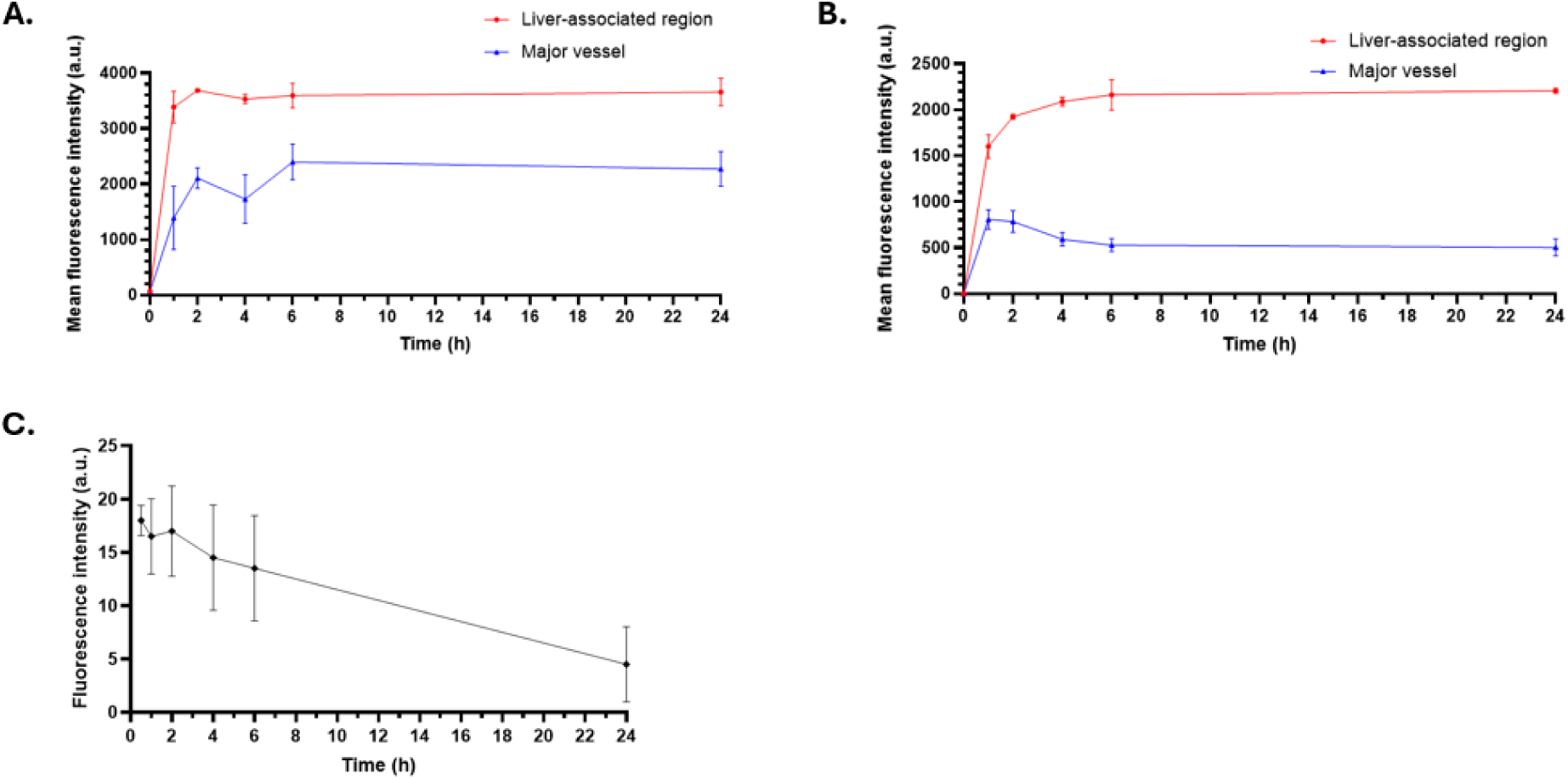
Quantification of *in vivo* and plasma fluorescence following intravenous administration of DiR-loaded PEGylated liposomes. Mean fluorescence intensity measured within liver-associated and major-vessel regions of interest over 24 h under NIR-I imaging conditions (A) and NIR-II imaging conditions (B) following intravenous administration of DiR-loaded liposomes using an *in vivo* imaging system. Plasma-associated fluorescence intensity measured using a plate reader at the indicated time points after injection (C). All results are mean ± SD (n = 2 mice).

### *Ex vivo* biodistribution of DiR-loaded liposomes 24 h after intravenous administration

Major organs were excised 24 h after injection and imaged under broadband, NIR-I, and NIR-II conditions (Figure 7A). Fluorescence accumulated preferentially in the liver and spleen, with only weak signal in kidneys, lungs, and heart, and none in organs from untreated controls. Quantification confirmed this pattern under both modalities (Figure 7B). Absolute intensities differed between the two acquisition settings, but the relative organ distribution was equivalent, indicating that NIR-II detection reports the tissue distribution of DiR-associated fluorescence similarly as with the established NIR-I window. The distribution itself (hepatic and splenic sequestration with negligible renal signal) is characteristic of PEGylated liposomes cleared by the mononuclear phagocyte system rather than eliminated renally.^18,19,22,23^

**Figure 7.**
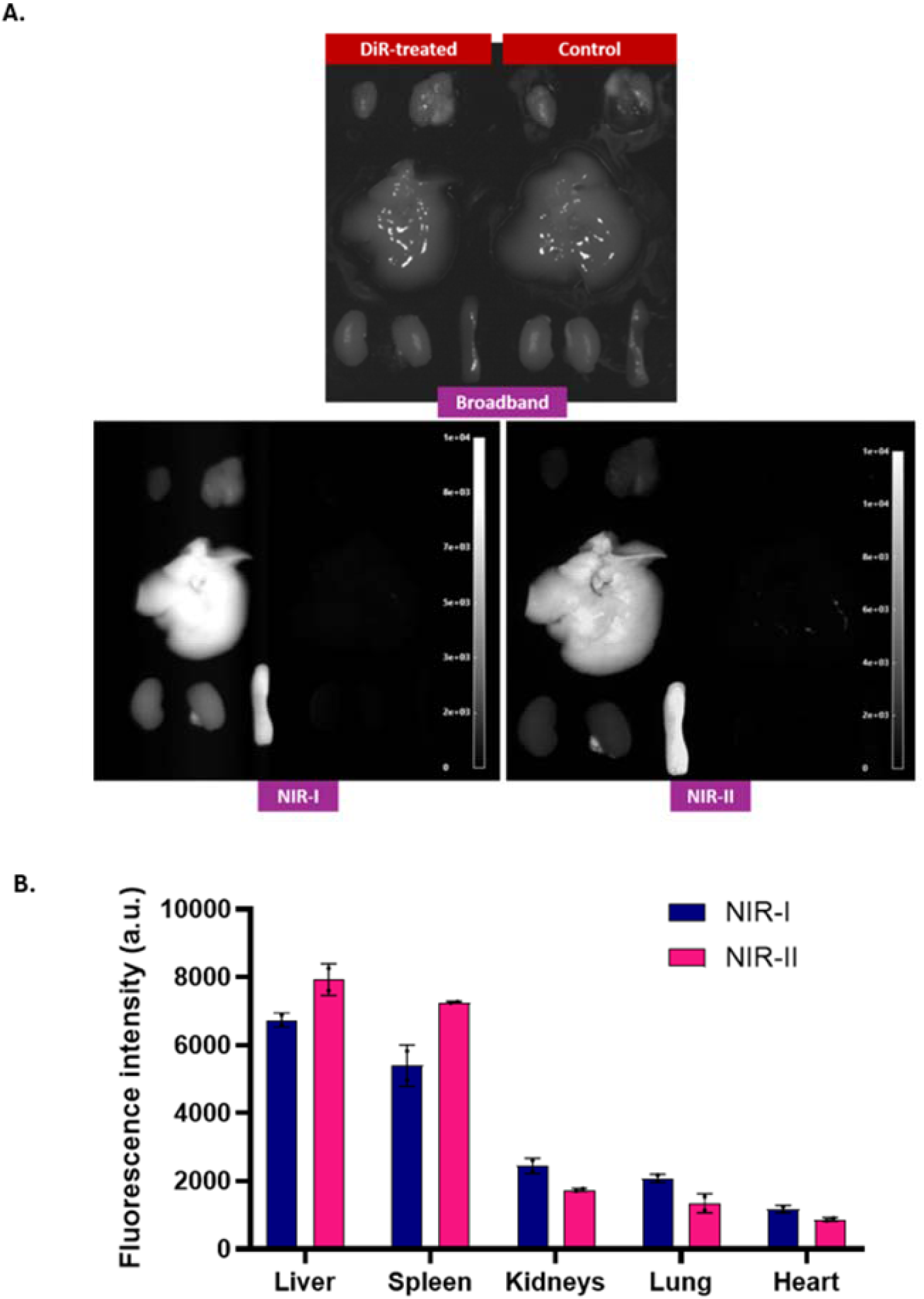
*Ex vivo* organ distribution of DiR-associated fluorescence 24 h after intravenous administration of DiR-loaded liposomes. Representative broadband, NIR-I, and NIR-II fluorescence images of major organs collected 24 h after intravenous administration of DiR-loaded liposomes, with corresponding organs from untreated control mice included for comparison (A). NIR-I images were acquired using 670 nm excitation and a BP785 emission filter, whereas NIR-II images were acquired using 760 nm excitation and a BP1400 emission filter. Fluorescence intensity measured in the liver, spleen, kidneys, lungs, and heart under NIR-I and NIR-II imaging conditions (B). All results are mean ± SD (n = 2 mice).

These *ex vivo* biodistribution results corroborate the *in vivo* imaging data and support the proposed screening and formulation workflow as a practical route to preclinical NIR-II imaging using a commercially available fluorophore.

## CONCLUSION

This work identifies commercially available fluorophores that exhibit NIR-II emission. Among the screened lipophilic dyes, the NIRF DiR was the only member of the DiO-DiA-DiI-DiD-DiR set to produce a strong signal in all twelve excitation-detection configurations; the far-red fluorophore DiD was detectable only under 670 nm excitation, and the visible fluorophores DiO, DiA, and DiI were indistinguishable from background throughout. Among water-soluble dyes, IRDye 680RD gave strong NIR-II fluorescence at 670 nm, S7 at 670 nm and 760 nm, and ICG at 808 nm excitation. NIR-II performance depended on fluorophore identity, and on excitation and emission wavelengths.

The hydrophobic NIRF DiR was selected for *in vivo* validation. It showed a linear concentration-response as a free dye under 760 nm excitation and BP1400 detection, retained long-wavelength fluorescence after insertion into PEGylated liposomes, and remained quantifiable within a defined linear range. Upon liposome injection in mice, NIR-II detection resolved vascular structures that NIR-I could not consistently delineate and yielded a circulating-phase time course paralleling the measured decline in plasma fluorescence. A biodistribution study exhibited similar liver- and spleen-dominated biodistribution profiles in the NIR-I and NIR-II imaging window. DiR is likely immediately applicable to NIR-II tracking of liposomes, lipid nanoparticles, extracellular vesicles, and labelled cell membranes, using the same passive incorporation already routine at NIR-I. Several of the water-soluble dyes are sold as NHS esters and other reactive derivatives, which would extend this approach to antibody, protein, and peptide labelling and to covalent tagging of nanoparticle surfaces, although the performance of the resulting conjugates will require validation.

In conclusion, NIR-II imaging is accessible to laboratories without synthetic chemistry capability, provided the fluorophore is carefully selected to match the available NIR-II excitation and emission configuration of the detection system.

## ACKNOWLEDGEMENTS

The authors gratefully acknowledge Dr. Nahyun Kwon (University of Toronto) for her valuable advice and critical review of the manuscript. S.M. gratefully acknowledges funding from Natural Sciences and Engineering Research Council of Canada (Discovery Grant RGPIN-2022-04384), Canada Foundation for Innovation (Fonds des leaders John-R.-Evans 42712). M.L.A.H. gratefully acknowledges funding from the Faculty of Pharmacy and ESP at Université de Montréal (Bourse de la Montagne) and a PhD scholarship from FRQNT.

## TOC

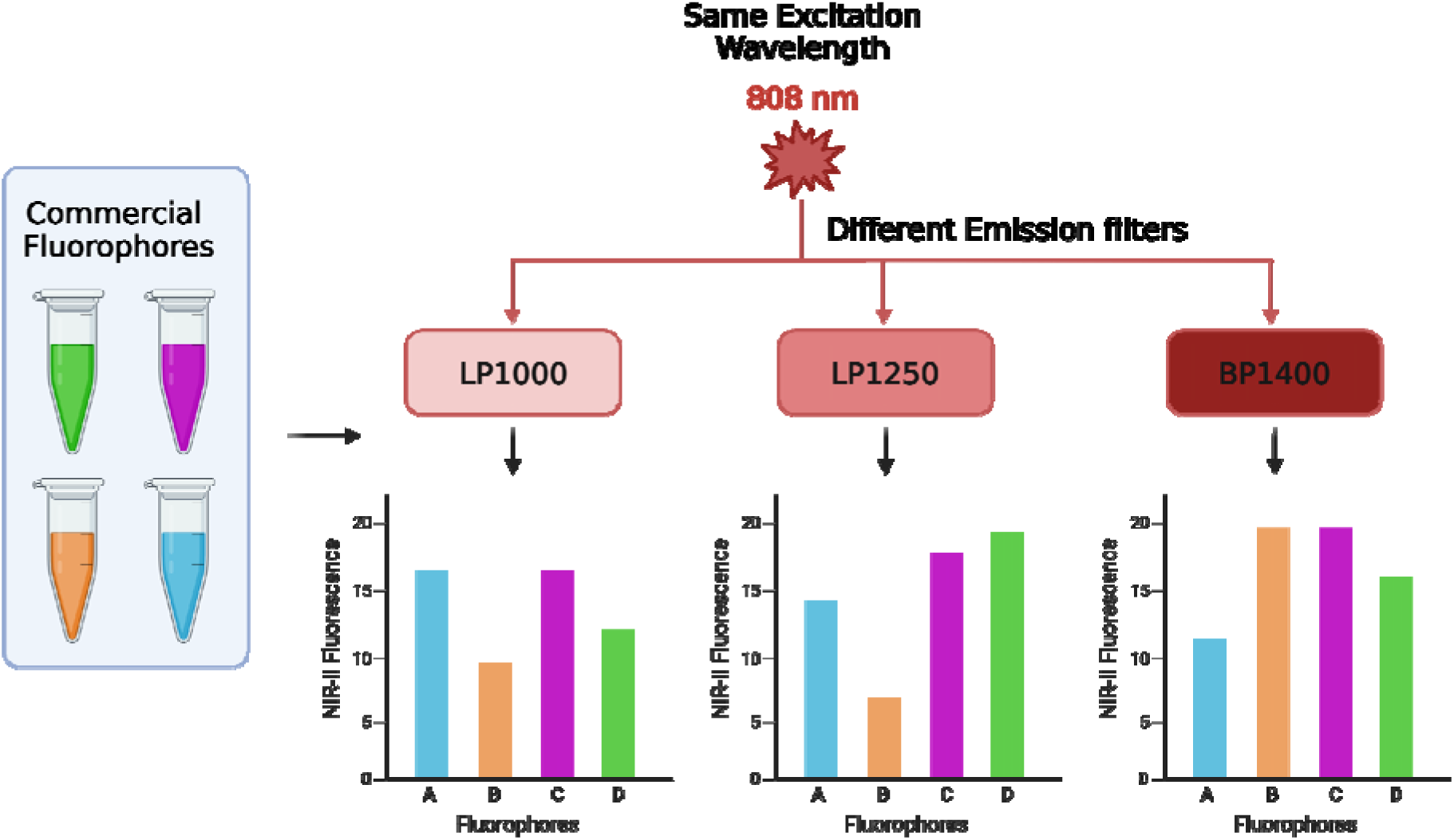

