## SUPPORTING INFORMATION (SI) for "Off-the-shelf NIR-I fluorophores as ready-to-use NIR-II probes: screening and in vivo validation"

Simon Matoori

Université de Montréal

2940 Chemin de Polytechnique, Montreal, QC H3T 1J4

**SUPPORTING INFORMATION (SI)**

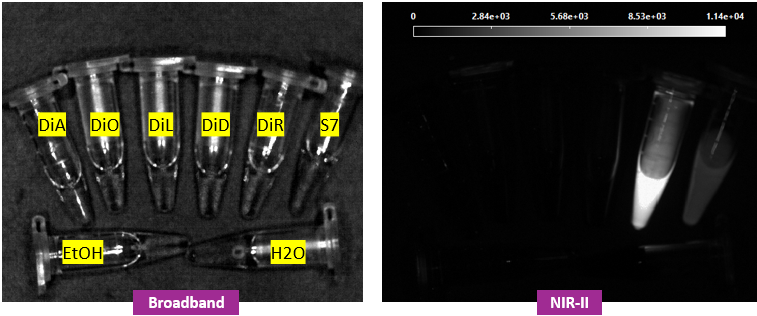

**Figure S1. Representative broadband and NIR-II fluorescence images of commercially available fluorophores.** Broadband images (A) and NIR-II fluorescence images acquired using 760 nm excitation and a BP1400 emission filter (B) of DiA, DiO, DiI, DiD, DiR, and Sulfo-Cyanine7 (S7) at a concentration of 0.5 µg/mL. Ethanol (EtOH) and ultrapure water (H₂O) were included as solvent controls.

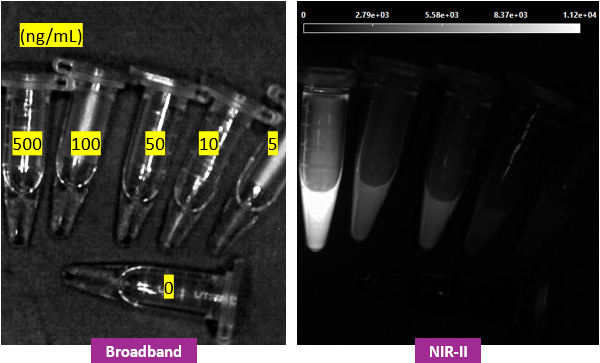

**Figure S2. Representative broadband and NIR-II fluorescence images of serial dilutions of free DiR.** Broadband images (A) and NIR-II fluorescence images acquired using 760 nm excitation and a BP1400 emission filter (B) of free DiR solutions at concentrations of 0, 5, 10, 50, 100, and 500 ng/mL in ethanol.

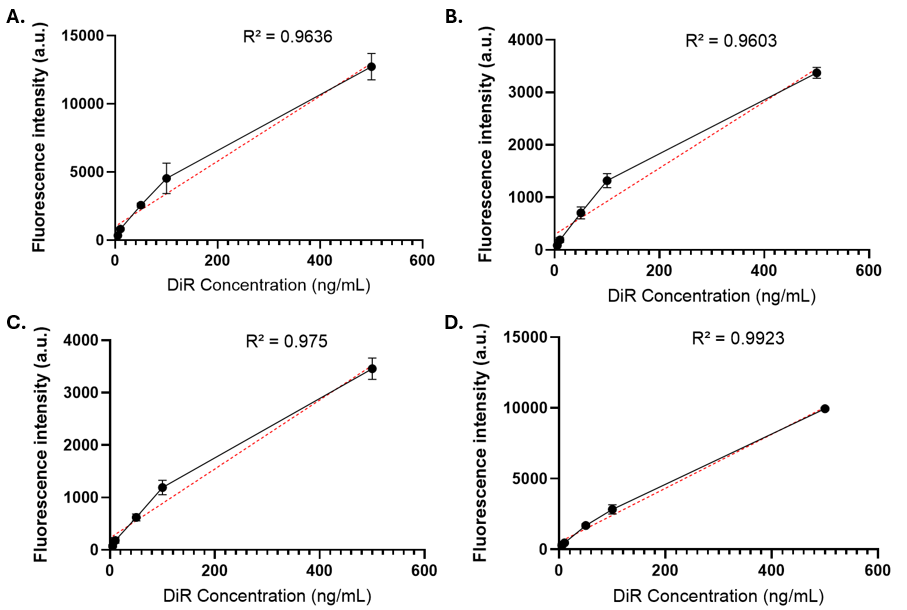

**Figure S3. Concentration-dependent fluorescence response of free DiR under 760 nm excitation using different emission filters.** Fluorescence intensity of free DiR solutions in ethanol at increasing concentrations using 760 nm excitation combined with a BP950 (A), LP1000 (B), LP1250 (C), or BP1400 (D) emission filter. Mean fluorescence intensity was determined within manually defined regions of interest and plotted as a function of DiR concentration. Individual measurements are shown as black points, and the red dashed lines represent the linear regressions. The corresponding coefficients of determination (R²) are indicated in each panel. All results are mean ± SD (n = 3).

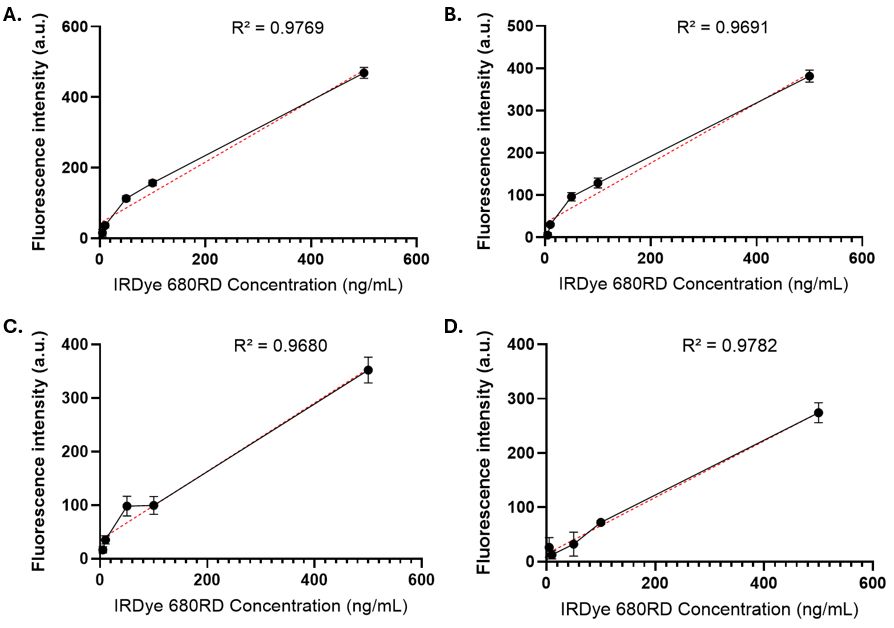

**Figure S4. Concentration-dependent fluorescence response of free IRDye 680RD under 670 nm excitation using different emission filters.** Fluorescence intensity of free IRDye 680RD solutions in water at increasing concentrations using 670 nm excitation combined with a BP950 (A), LP1000 (B), LP1250 (C), or BP1400 (D) emission filter. Mean fluorescence intensity was determined within manually defined regions of interest and plotted as a function of IRDye 680RD concentration. Individual measurements are shown as black points, and the red dashed lines represent the linear regressions. The corresponding coefficients of determination (R²) are indicated in each panel. All results are mean ± SD (n = 3).

| **Fluorophore** | **Molar mass (g/mol)** | **Concentration (µmol/L)** |
| --- | --- | --- |
| **DiO** | 782.27 | **0.639 µM** |
| **DiA** | 787.06 | **0.635 µM** |
| **DiL** | 961.34 | **0.520 µM** |
| **DiD** | 987.38 | **0.506 µM** |
| **Alexa Fluor 647** | 959.26 | **0.521 µM** |
| **IRDye 680RD** | 1003.46 | **0.498 µM** |
| **DiR** | 1013.42 | **0.493 µM** |
| **Sulfo-Cyanine7** | 844.05 | **0.592 µM** |
| **IRDye 800CW** | 1166.2 | **0.429 µM** |
| **ICG** | 774.96 | **0.645 µM** |
| **Sulfo-Cyanine7.5** | 1180.47 | **0.424 µM** |

**Table S1. Molar concentrations of commercially available fluorophores used for screening.** Molar concentrations were calculated from the molecular mass of each fluorophore at a fixed mass concentration of 0.5 µg/mL, equivalent to 0.5 mg/L. Owing to differences in molecular mass, the resulting molar concentrations ranged from 0.424 to 0.645 µmol/L. Screening experiments were performed at equal mass concentrations.
